# Phyllochron duration and changes through rice development shape the vertical leaf size profile

**DOI:** 10.1101/2022.03.12.484079

**Authors:** Janet P. Pablo, Benoit Clerget, Crisanta Bueno, Jacqueline Dionora, Abigail J. Domingo, Constancio C. De Guzman, Edna A. Aguilar, Nina M. Cadiz, Pompe C. Sta Cruz

## Abstract

- Well-irrigated aerobic rice (*Oryza sativa* L.) plants are often shorter with smaller leaves, and the leaf appearance rate is slower in well-irrigated than in flooded environments. This observation questions the functional relationship between the leaf appearance rate, which is correlated with the apical cell production rate, and leaf length that is in turn correlated with the leaf division zone length.
- Meristem size, blade size, and appearance were monitored for all leaves from leaf 6 on the main stem of rice plants in field experiments using two varieties, two watering systems, and three sowing dates.
- Leaf blade and division zone lengths were correlated in all leaves of the main stem of rice plants. New stable linear relationships were found between the leaf division zone growth duration and internal cell production rate, which changed after leaf 11. These stable relationships imply a stable link between the rates of cell production in the stem apex and leaf meristem, explaining the vertical leaf size profile.
- Overall, faster leaf appearance leads to shorter leaves and organs in rice because of a functional link, which is counterbalanced by additional acclimation in transplanted plants.

## INTRODUCTION

In cereals, leaf primordia are initiated by the apical bud, with a stable thermal time interval called the plastochron, whereas leaf blades emerge from the whorls of the preceding sheaths at stable thermal time intervals called phyllochrons (Gallagher, 1979; Wilhelm & McMaster, 1995). The phyllochron can remain stable throughout plant development, assume different successive values, or evolve continuously. In rice (*Oryza sativa* L.), three successively increasing phyllochron durations have been reported, with a faster appearance of the first leaves already present in the embryo and a slower appearance of the last four leaves (Yoshida, 1981; Nemoto *et al*., 1995).

The leaf kinetics of rice differ from those of other cereals. Specifically, rice plastochrons and phyllochrons are equal, such that the observed number of leaves growing inside the sheaths is always four (Nemoto & Yamazaki, 1993; Nemoto *et al*., 1995). Thus, in rice, it takes four phyllochrons for leaf development and growth between primordium initiation and tip appearance and another phyllochron to complete the development and growth of the blade and sheath. In other words, the leaf initiation and appearance rates of rice change simultaneously.

The phyllochron is primarily sensitive to temperature, and leaf appearance is often linear when plotted against the thermal time of the shoot apex (Wilhelm & McMaster, 1995; Vinocur & Ritchie, 2001). In addition, many other environmental factors, such as nutrient or water availability, light, and CO_2_ concentration, affect phyllochrons, together with plant morphology. In cereals, development and growth are restricted to a specific zone at the base of the growing leaf. Initially, simultaneous anticlinal divisions (perpendicular to the plane of the layer) occur in two epidermal layers in a small area at the base of the stem apical dome (Chonan, 1993; Maeda & Miyake, 1993). Subsequently, a few (often 3–5 in a rank in longitudinal sections) periclinal divisions (parallel to the plane) of the contiguous cells of both layers occur, creating a small protuberance of approximately 30 µm in diameter—the primordium (Itoh *et al*., 1998). The primordium diameter is one-third to half of the apical dome height (50–70 µm), which is covered by lines of 14–15 epidermal cells at the time of appearance of leaves 7–12. Similar observations have reported for the primordia of leaves 11 and 12 in apical longitudinal sections of IR 64 (de Raissac, CIRAD, Montpellier, France, unpublished data). In the newly generated division zone, all undifferentiated cells multiply actively, and the length of the meristematic blade exponentially increases during this phase (Ben-Haj-Salah & Tardieu, 1995; Lafarge *et al*., 1998). The cells are organised in parallel lines and permanently pushed out by younger developing cells. In fescue leaf 4, wheat leaves 4–11, and rice leaves 6–7, some distal cells of the three-phyllochron-old leaves enter the elongation zone simultaneously with the tip appearance of the previous leaf (Fournier *et al*., 2005; Parent *et al*., 2009). In this new zone, the cells elongate to their maximum length without further division. In rice, cell flow to the elongation zone rapidly increases and peaks half a phyllochron later (Parent *et al*., 2009). The length of the division zone remains stable during one phyllochron, because the cell flow rate to the elongation zone is equal to the production rate of new cells in the division zone. Leaf blades elongate linearly during this steady-state period (Schnyder *et al*., 1990; Ben-Haj-Salah & Tardieu, 1995; Fiorani *et al*., 2000; Parent *et al*., 2009). In rice, this period lasts for approximatively one phyllochron. Finally, elongated cells enter the maturation zone, which is characterised by secondary cell wall deposition and chloroplast formation. The elongation zone reaches the maximum length at the tip appearance. Probably triggered by the leaf-tip appearance, division activity is gradually transferred from the blade to sheath, and the collar zone passively moves through the division zone (Skinner & Nelson, 1995), simultaneous with ligule differentiation. Collar emergence from the enclosing sheath probably triggers the end of sheath development, followed by the consecutive onset of internode development (Fournier *et al*., 2005).

In cereals, leaf length increases with leaf rank up to the maximum size and decreases with the last rank, forming a bell-shaped profile (Elings, 2000; Dornbusch *et al*., 2011). In wheat, the length of the leaf-growing zone and the rate of leaf elongation linearly increase with leaf rank (Kemp, 1980). Similarly, in four *Poa* species with contrasting leaf lengths, longer leaves of rank 7 had longer division and elongation zones and faster leaf elongation rates (Fiorani *et al*., 2000). In contrast, the lengths of the division zone and blades are negatively correlated with the rate of leaf appearance. In rice *plastochron1* and *plastochron2* mutants, the rate of leaf appearance was doubled simultaneously with the halving of the leaf and internode lengths because of fewer cells of unchanged size (Kawakatsu *et al*., 2006). Similar negative correlations have been noted between the rate of leaf appearance and either the size of leaf 7 in 190 rice varieties (Rebolledo *et al*., 2012) or the sizes of leaves 1 and 2 in two sets of 28 and 50 wheat varieties (Rebetzke & Richards, 1999). Finally, in chrysanthemum and rice, shorter plastochrons are correlated with a greater rate of cell division in the apical meristem and larger apical meristem sizes (Nougarède *et al*., 1990; Itoh *et al*., 1998). In chrysanthemum, the duration of cell number doubling was shorter, although the duration of mitosis itself did not vary.

Well-irrigated aerobic rice crops typically yield less than flooded crops. Moreover, well-irrigated aerobic rice plants are smaller (leaves, stems, and panicles) and show a lower rate of leaf appearance, leading to fewer leaves on each stem (Lafitte *et al*., 2002; Bouman *et al*., 2007; Clerget *et al*., 2014). Thus, the possible link between plant size and leaf appearance rate was speculated. With contrasting phyllochrons before and after the appearance of the 12^th^ leaf, no correlation was found between the rate of leaf appearance and the rate of leaf elongation (Egle *et al*., 2015).

To this end, the present study aimed to evaluate the relationship between the variable rate of leaf appearance and the vertical profile of leaf blade length. Leaf blade length, division zone length, and cell count per file in the division zone of leaf intercalary meristems were monitored in three successive field experiments involving two rice varieties and two watering systems.

## MATERIALS AND METHODS

### Experimental design and crop management

Experiments were conducted at the IRRI research farm in Los Baños, Laguna, Philippines (14°11′N 121°15′E, 21 m a.s.l.) from January to December 2015. For three separate experiments, seeds were sown on 22 and 26 January 2015, 1 June 2015, and 11 September 2015 in 11 × 1.5 m plots. The rice variety IR 72—an IRRI-improved variety maturing in 110–120 days—was used during the three sowing months. In June, a second variety, SIAM 29, a Malaysian landrace that is highly photoperiod-sensitive even at latitudes close to the equator (Dore, 1959), was added. This is a late variety when sown on growing days and produces many leaves during the long vegetative phase, which is expected to last until the autumn equinox (Clerget *et al*., 2021). Seeds were obtained from the IRRI rice Genebank. The crops were either flooded with 3–5 cm of water from 3 weeks after sowing up to 2 weeks before harvest or grown under aerobic conditions with the soil maintained close to water saturation via gravity irrigation triggered when the water potential at 15 cm reached -10 kPa. The seeds were soaked for 24 h and then incubated at room temperature (daily mean of 27–29°C) for 24–48 hours prior to sowing. Pre-germinated seeds were manually sown at a hill spacing of 20 × 20 cm, with one seedling per hill. Basal fertilisation with single superphosphate (40 kg P ha^-1^), muriate of potash (40 kg K ha^-1^), and zinc sulphate (10 kg Zn ha^-1^) was applied before sowing. Urea was applied (40 kg N ha^-1^ per application) in three and five splits for IR 72 under flooded and aerobic conditions, respectively. Additional applications were performed every 3 weeks for Siam 29. Weeds were manually removed, and pests were controlled using the recommended chemicals to ensure that the plant samples were healthy.

### Measurements

#### Leaf appearance rate and leaf size

Leaves on the main tillers of consecutive plants from 3 to 6 lines were regularly tagged with their rank of appearance. Leaves were numbered acropetally (McMaster, 2005), with the prophyll tagged as #1. The leaf appearance of five consecutive plants was recorded three times a week using Haun leaf decimal scale units (Haun, 1973). In rice, the appearance of the collar of leaf N (when Haun value = N) and the tip of the leaf N+1 are synchronous. Accordingly, the number of already appearing leaves (LN) was estimated by adding one to the Haun values. Using the same plants, leaf length was measured from the collar to the tip of each fully expanded leaf, whereas leaf width was measured at the widest part of the leaf. Leaf area was computed as the product of the length and width multiplied by 0.72.

#### Size of the division zone in leaf intercalary meristem

Shortly before the appearance of each leaf from rank six to the flag-leaf, five purposively selected plants were collected before the appearance of the targeted leaf tip (Haun stage ≈ N.7, with N+1 being the target leaf rank). The next leaf to appear, still growing within the previous leaf sheaths, was then in the steady phase of development, and the length of the meristem division zone was thus maximum and stable (Parent *et al*., 2009). The next leaf to appear in each selected plant was excised at the point of attachment to the node and its length was recorded (Supporting Information Fig. S1). Approximately 10 mm at the base of the developing leaf was cut off and subjected to sequential tissue clearing using solutions of ethanol (95%), sodium hydroxide (5%), and sodium hypochlorite (20%). This clearing method was selected because of the better clarity of images and shorter processing time. Prior to clearing, leaf tissues were fixed in 2.5% glutaraldehyde, allowed to sink into a fixative solution using a desiccator, repeatedly washed with 50 mM phosphate buffer or Nanopure water, and stored in 70% or 95% ethanol before further clearing. The coiled leaves were then slowly unrolled using dissecting needles under a 2× magnifier (Carson Deskbrite 200, Carson Optical, USA). Adaxial leaf tissues were imaged using a microscope (Olympus BX61, Olympus Corporation, Japan). Montages of the leaf blade meristem were constructed from photomicrographs of successive image segments to form an entire strip from the base to the end of the leaf blade epidermal division zone. Three types of cell files were observed in the epidermal division zone: two to three intercostal files of short and narrow cells alternating with either four to five files of longer and wider cells covering the prominent part of the veins or two to three files of bulliform cells covering the depressed area between two veins (Supporting Information Fig. S2A&B). The length of the division zone was measured from the leaf base to the first occurrence of asymmetrical formative divisions yielding two stomatal initial cells of very unequal length, which signalled the edge of the proliferative division zone in both stomatal and adjacent intercostal cell lines (Fiorani *et al*., 2000) (Supporting Information Fig. S3). The fate of the pairs of asymmetrical cells was assessed during experiment 1. For instance, in leaf 10 of the main stem of an IR 72 plant, the first asymmetrical divisions were observed 4.1 mm from the leaf base at the distal end of a file of 609 undifferentiated cells (Supporting Information Fig. S4A). From 4.1 mm, the proportion of half-moon cells increased (B) up to 50% at 5.5 mm, where all cells in the stomatal files were paired (C). No immature cell walls other than the half-moon shaped walls were observed in the 4.1–5.5 mm zone, where only formative divisions were observed. Between 5.5 and 5.6 mm, the length of the half-moon cells doubled and both paired cells further differentiated (D), as reported by Luo *et al*. (2012).

Consecutive cell walls in the entire division zone were tagged along a file of future stomatal cells per leaf using ImageJ (NIH, USA).The number of cells in the file, and the length and position of each cell were recorded.

#### Climatic data

Climatic data for each experiment were recorded using a weather station installed at one end of the field. The incident global solar radiation (either GS1 dome solarimeter, Delta-TDevices, Cambridge, UK, or LI-200 Pyranometer, Li-Cor, Lincoln, NE, USA), air temperature at 2 m above the ground level (HMP45C, Vaisala, Helsinki, Finland), and soil temperature at a depth of 2 cm (T thermocouples, PyroControle, Vaulx en Velin, France) were measured at 1 min intervals, averaged on an hourly basis, and stored in data loggers (CR1000, Campbell Scientific, Logan, UT, USA). In the aerobic experiment, soil water tension at a depth of 15 cm was recorded daily using three tensiometers (Jet Fill 2725, Soil Moisture Equipment Corp., Goleta, CA, USA).

### Data analysis

#### Sum of temperatures

Thermal time was calculated on an hourly basis using the soil temperature up to the onset of stem elongation (as an estimate of shoot apex temperature), after which air temperature was used. Cardinal temperatures were assumed to be 11°C for the base temperature (Lafarge *et al*., 2010) and 30°C and 42°C for the optimum and maximum temperatures, respectively (Yin *et al*., 1995). Linearity of response was assumed between the cardinal temperatures, with the progress of thermal time equal to zero at temperatures below the base or above the maximum.

#### Leaf appearance kinetics

The leaf appearance rate was assumed to follow a trilinear broken-stick model in which the first leaves appeared faster than the succeeding leaves. The observed leaf number on the main stem (*LN*) was regressed against the elapsed thermal time from plant emergence (*TT*) using the following equation:

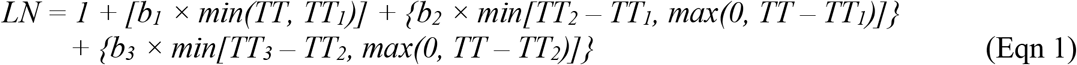

where the number of leaves at emergence was assumed to be 1; *b*_*1*_, *b*_*2*_, and *b*_*3*_ are the initial, secondary, and tertiary rates of development, respectively; *TT*_*3*_ is the thermal time at which the last leaf appeared; and *TT*_*1*_ and *TT*_*2*_ are the thermal times at which the development rate changed. The parameters and their confidence intervals were iteratively estimated using the NLIN procedure in SAS (2012). *Phyllochron1, phyllochron2*, and *phyllochron3* were calculated as the inverses of *b*_*1*_, *b*_*2*_, and *b*_*3*_, respectively. For Siam 29, a quadrilinear model was used because an additional broken segment was required to fit the end phase of the observed leaf appearance.

As reported by Egle *et al*. (2015), *phyllochron2* gradually increases to *phyllochron3* from the appearance of leaf_*TT2-1*_ to the appearance of leaf_*TT2+1*_, where leaf_*TT2*_ is the leaf appearing at the breaking point *TT*_*2*_ (Eqn 1). To increase the precision of the phyllochron_i_ value at the time of each leaf_i_ appearance, the phyllochron_i_ values from leaf_*TT2-1*_ to leaf_*TT2+1*_ appearances were computed as follows:

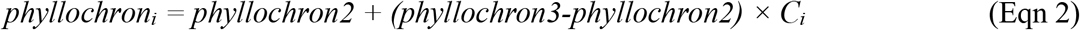

where *C*_*i*_ is a coefficient with values of 0.25, 0.5, and 0.75 for leaf_*TT2-1*_, leaf_*TT2*_, and leaf_*TT2+1*_, respectively.

The length of the leaf division zone no longer increases from the half-phyllochron before leaf tip appearance (i.e. 3.5 phyllochrons after leaf initiation) (Fournier *et al*., 2005; Parent *et al*., 2009). However, since we only measured the proliferative division zone in the present study, we assumed that the actual action of the synchronising cue received at the tip appearance of the previous leaf would trigger the switch from proliferative to formative division at the distal end of the still homogeneous meristematic young leaf. Consequently, the growth of the proliferative division zone would end with phyllochron3, and the formative division zone would grow until 3.5 phyllochrons.

For each leaf, the duration of three phyllochrons from initiation (*P*_*3*_) was computed as follows:

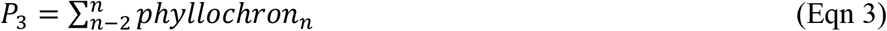

where *phyllochron*_*n*_ is the phyllochron value at the appearance of leaf_n_.

### Modelling of epidermal division zone size in response to its developmental duration

In the absence of further information, a stable mean rate of cell doubling duration was assumed during the initial three phyllochrons of the lifespan of each leaf. Each epidermal cell file was assumed to originate from an initial line of three consecutive synchronised cells from the tunica layer, as observed in the rice embryo (Maeda and Miyake 1993) up to leaf 12 (Maeda & Miyake, 1993; Itoh *et al*., 1998, 2005) (Supporting Information Fig. S5). These three cells generated both the adaxial and abaxial epidermal cell files. Thus, the cell number in the longest lines of the adaxial epidermal division zone followed an exponential increase of two to the power of the average number of multiplication cycles from leaf primordium initiation until tip appearance of the preceding leaf (i.e. for the duration of three phyllochrons) (Fiorani & Beemster, 2006). Addressing epidermal cells simplified the rationale, as this tissue arose from the apical meristem epidermis initially covering the leaf primordium. For each leaf, after a number of division cycles (*nb_cycles*_*P3*_) occurred during three phyllochrons, the number of cells issued from 1.5 initial leaves in the division zone of the abaxial epidermis (*cell_nb*_*P3*_) was computed as follows:

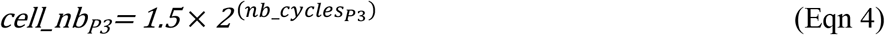

Analyses of the variance of leaf blade and leaf division zone sizes and parameters were carried out using the GLM procedure ii SAS (2012), assuming sowing date, variety, watering, and leaf rank as fixed factors. Linear regressions between traits were computed using the REG procedure in SAS (2012).

## RESULTS

### Climatic data

The mean air temperature was higher during June sowing (29.5°C) than during January (25.3°C) and September (26.3°C) sowing (Supporting Information Fig. S6A). The monthly mean soil temperature was higher than the mean air temperature in the first month of cultivation for all sowing dates, as the daily maximum soil temperature was much higher than the maximum air temperatures, particularly during June and September sowing. During the first cropping month, the plants did not shade the soil; therefore, the soil received total solar radiation.

The mean daily solar radiation was the same during January (18.4 MJ m^-2^) and June (18.4 MJ m^-2^) sowings, but showed a markedly different distribution over the cropping season (Supporting Information Fig. S6B). In the last cropping month, when the grains were filled, the mean daily solar radiation was respectively 22 and 17.5 MJ m^-2^ for January and June sowings. For the September sowing, the mean daily solar radiation was 14.4 MJ m^-2^.

### Leaf appearance rate and leaf development duration

Despite large differences in origin and phenology, the leaf appearance kinetics of the late variety Siam 29 were very similar to those of IR 72. A quadrilinear model was used to fit the kinetics of Siam 29 leaf appearance, which showed three breaking points (Fig. 1A and Table 1). The third slowdown was observed after the appearance of the 21^st^ leaf at 1813°Cd or 103 days after sowing. The phyllochron changed gradually from leaves 11 to 14 over 2 weeks (Supporting Information Fig. S7). The third phase of this kinetics lasted for 1274°Cd or 74 days and was remarkably linear. The last phyllochron was very long (275°Cd or ∼17 days).

**Fig. 1:**
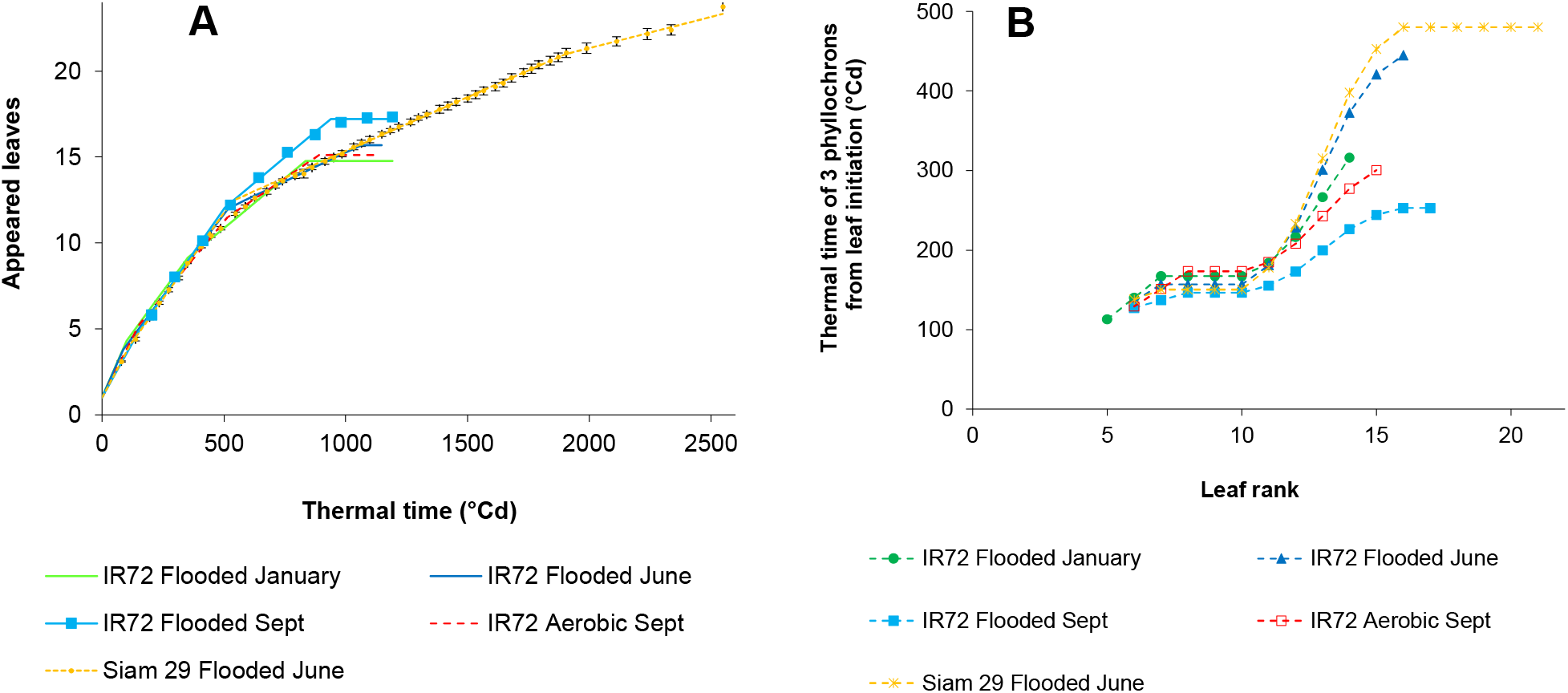
**A)** Observed (symbols) and predicted (lines) numbers of appeared leaves on the main stem with thermal time. Error bars indicate the 95% confidence interval of each mean value for the variety Siam 29. **B)** Estimated duration of three phyllochrons from the initiation of each leaf.

**Table 1:**
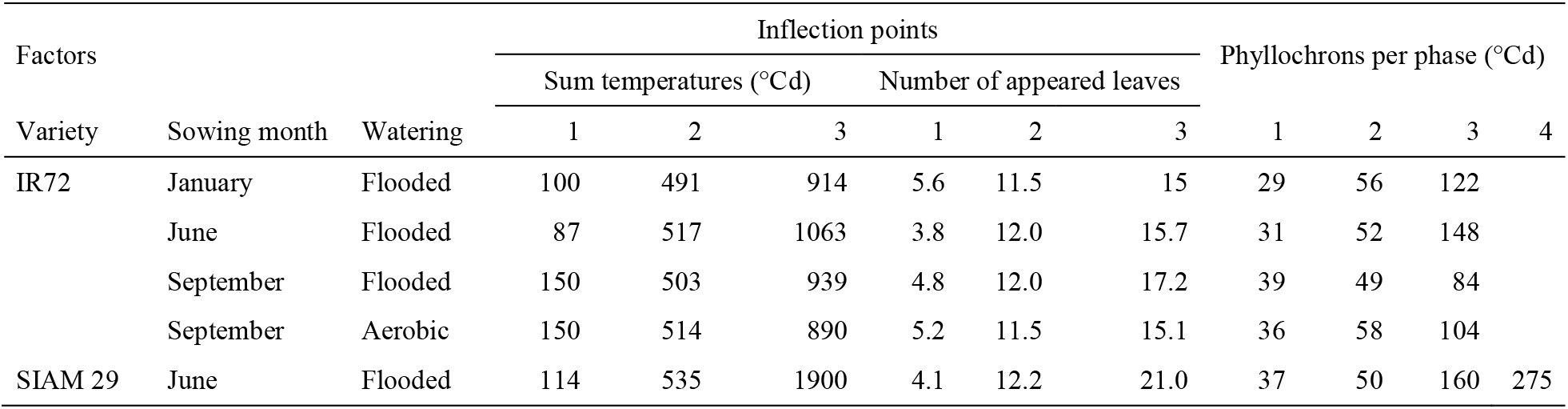
Parameters of the trilinear and quadrilinear models of the leaf appearance kinetics in the three experiments.

All leaf appearance kinetics of IR 72 fit the trilinear model well (Fig. 1A and Table 1). The upper leaves of the flooded IR 72 plants sown in September appeared at a faster rate (84°Cd leaf^-1^) than those of the plants sown in January and June (122°Cd and 148°Cd leaf^-1^, respectively). Meanwhile, the upper leaves of plants sown in September under aerobic conditions appeared at a slower rate (104°Cd leaf^-1^) than those of plants sown under flooded conditions.

The increases in phyllochron duration, approximately at the time when the 5^th^ and 12^th^ leaves appeared, prolonged the division zone growth duration (Fig. 1B). The first change affected three developing leaves, from 5 to 7, and the second gradually affected five leaves, from 11 to 16 leaves.

### Vertical leaf size profile

Regardless of rank, Siam 29 leaves were significantly longer, wider, and larger than IR 72 leaves (Fig. 2A,B,&C and Table 2). The maximum leaf length and width were reached for leaf 13, after which leaf sizes plateaued to leaf 16. The vertical profiles were incomplete in Siam 29. At the end of the experiment on 27 October (5 months after sowing), 21 leaves were entirely grown, five were developing, and panicle initiation had not yet occurred.

**Fig. 2:**
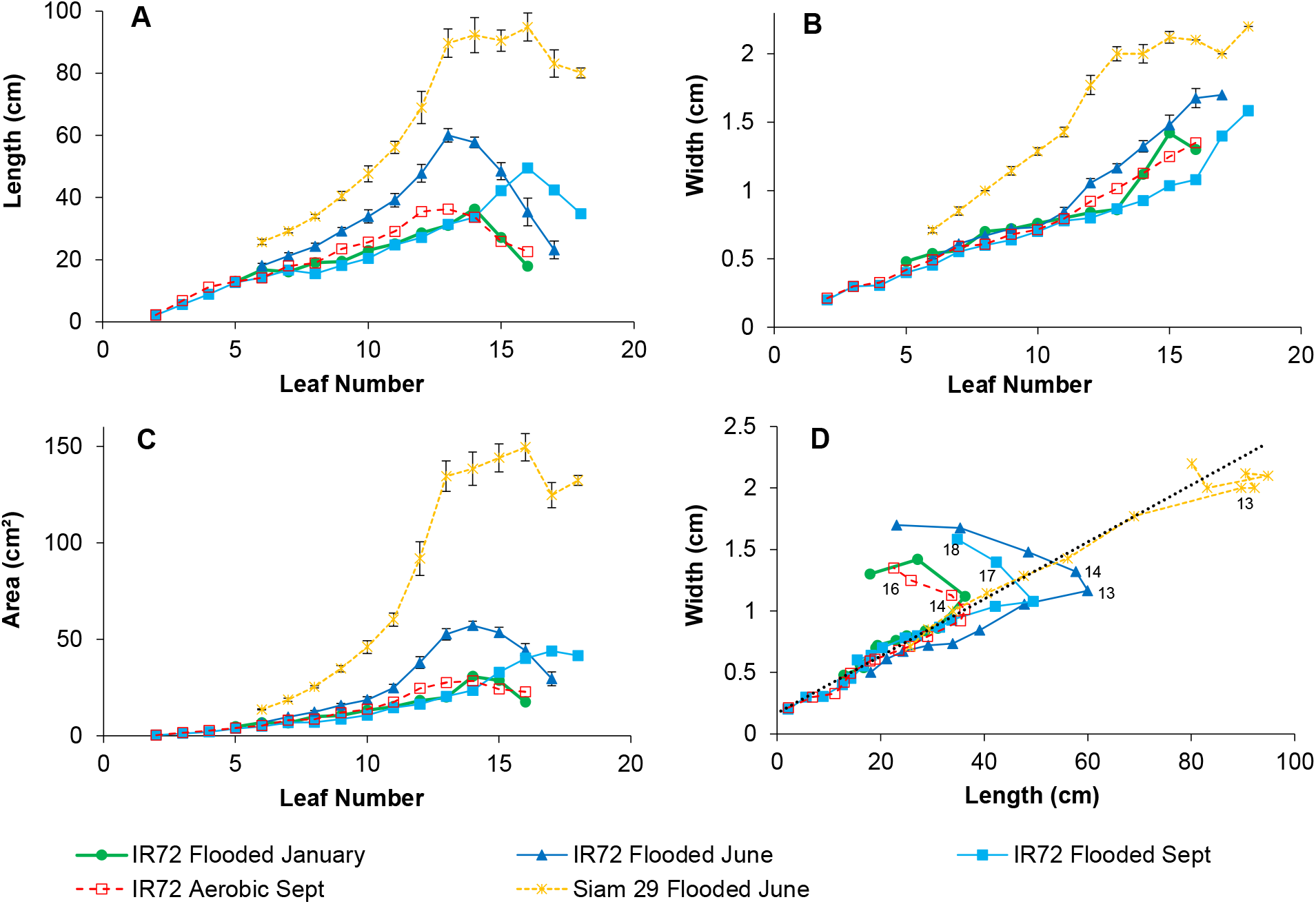
Leaf blade sizes by leaf rank on the main stem for the three experiments: mean length (A) and width (B), computed mean area (C), and relationship between the mean length and width (D). Error bars indicate the 95% confidence interval, of each mean value for the IR 72 and Siam 29 varieties sown on June. The black dashed line in D is drawn from the regression on leaves 6 to 12 of Siam 29. Rank of some leaves is tagged.

**Table 2:**
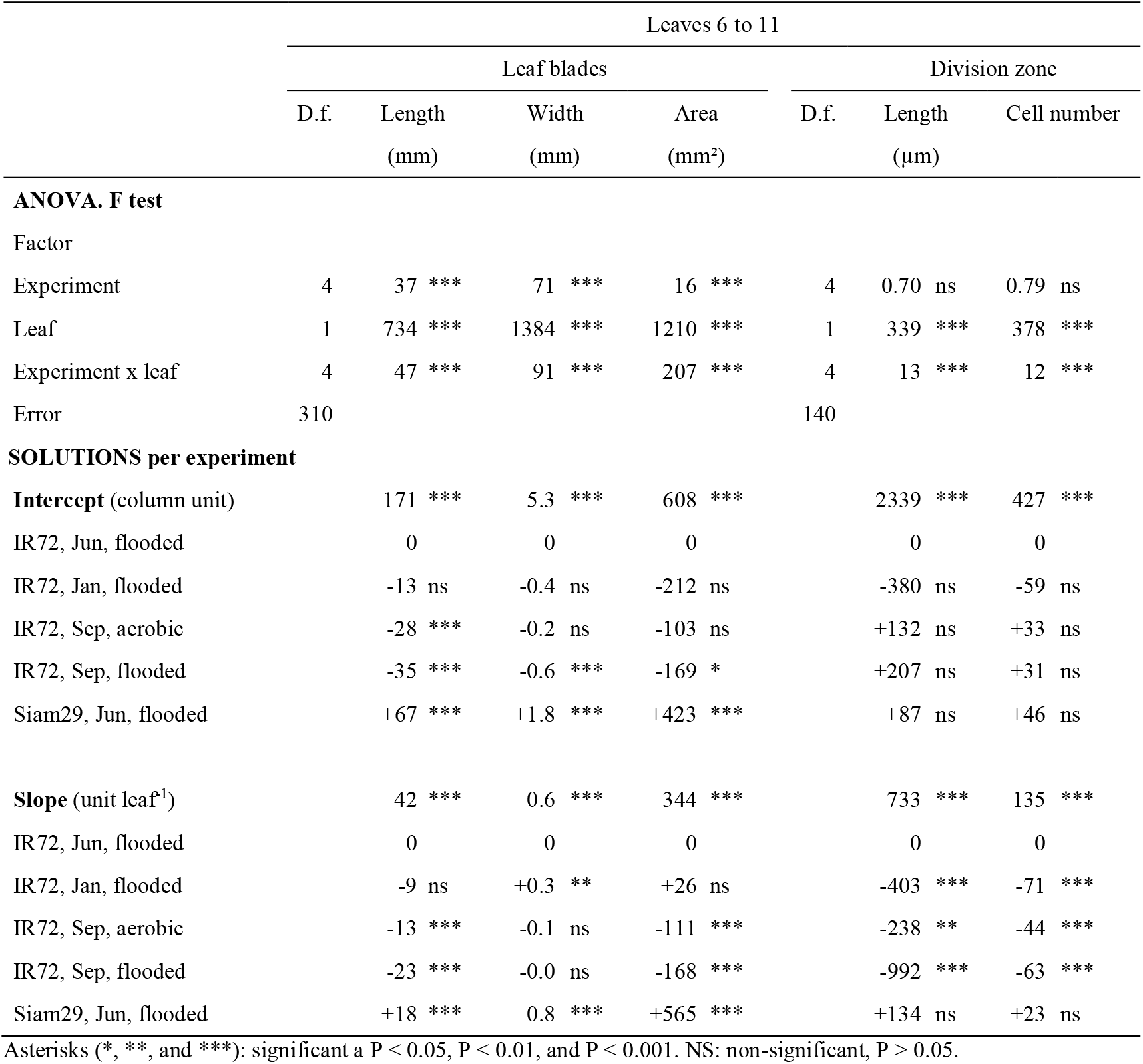
ANOVA and parameter estimates of the linear model for five traits increasing linearly with leaf rank in leaves 6 to 11.

The vertical leaf size profiles of IR 72 plants grown under flooded conditions were significantly modified by the sowing date and watering method. All leaves over rank 8 of IR 72 plants sown in June were longer and wider than their counterparts sown in January and September. Leaves from ranks 8 to 14 of IR 72 plants sown in September and grown under aerobic conditions were significantly larger than those of plants grown under flooded conditions.

The length and width of leaves up to rank 12 were linearly correlated (R^2^= 0.86, P<0.0001 for IR 72; R^2^= 0.95, P<0.0001 for Siam 29) (Fig. 2D). The larger leaves above rank 10–12 had a slightly narrower shape, whereas the last 2-3 leaves were shorter and wider than the previous leaves.

### Vertical profile of epidermal leaf meristem division zone length

Over twenty thousand 170-μm-long photomicrographs were stitched into 253 pictures of an entire strip of 253 leaf division zones. The half-moon shape of small new cells resulting from asymmetrical formative division clearly signalled the end of the proliferative division zone in the stomata files, as described by Fiorani *et al*. (2000) (Supporting Information Figs. S3 and S4).

Small half-moon cells are the precursors of the stomatal mother guard cells (Luo *et al*., 2012). Occasional asymmetrical divisions were also observed in the files of sister cells adjacent to a stomatal row, starting at the same distance from the leaf base. Masle (2000) reported that small cells were the precursors of trichomes in the corresponding cell files in wheat. No proliferative division with straight immature cell walls was observed more distally than the point where the first asymmetrical division occurred in both types of cell lines.

The length of the epidermal leaf meristematic division zone increased with leaf rank from 3 (6^th^ leaf) to 9.3 mm for leaf 14 in IR 72 and to 13.3 mm for leaf 16 in Siam 29 and then plateaued for leaves of the upper ranks (Fig. 3A). The vertical profiles of cell number per file and division zone length of the two varieties followed similar patterns, with Siam 29 showing significantly higher rates of increase with leaf rank (Fig. 3A&B, and Table 2). In all leaves, the 20-cell moving average of cell length asymptotically decreased from to 10–15 µm near the leaf base to 4 µm at the distal end of the division zone (Supporting Information Fig. S8). This decrease was discontinuous, showing alternate zones with either longer or shorter mean cell lengths.

**Fig. 3:**
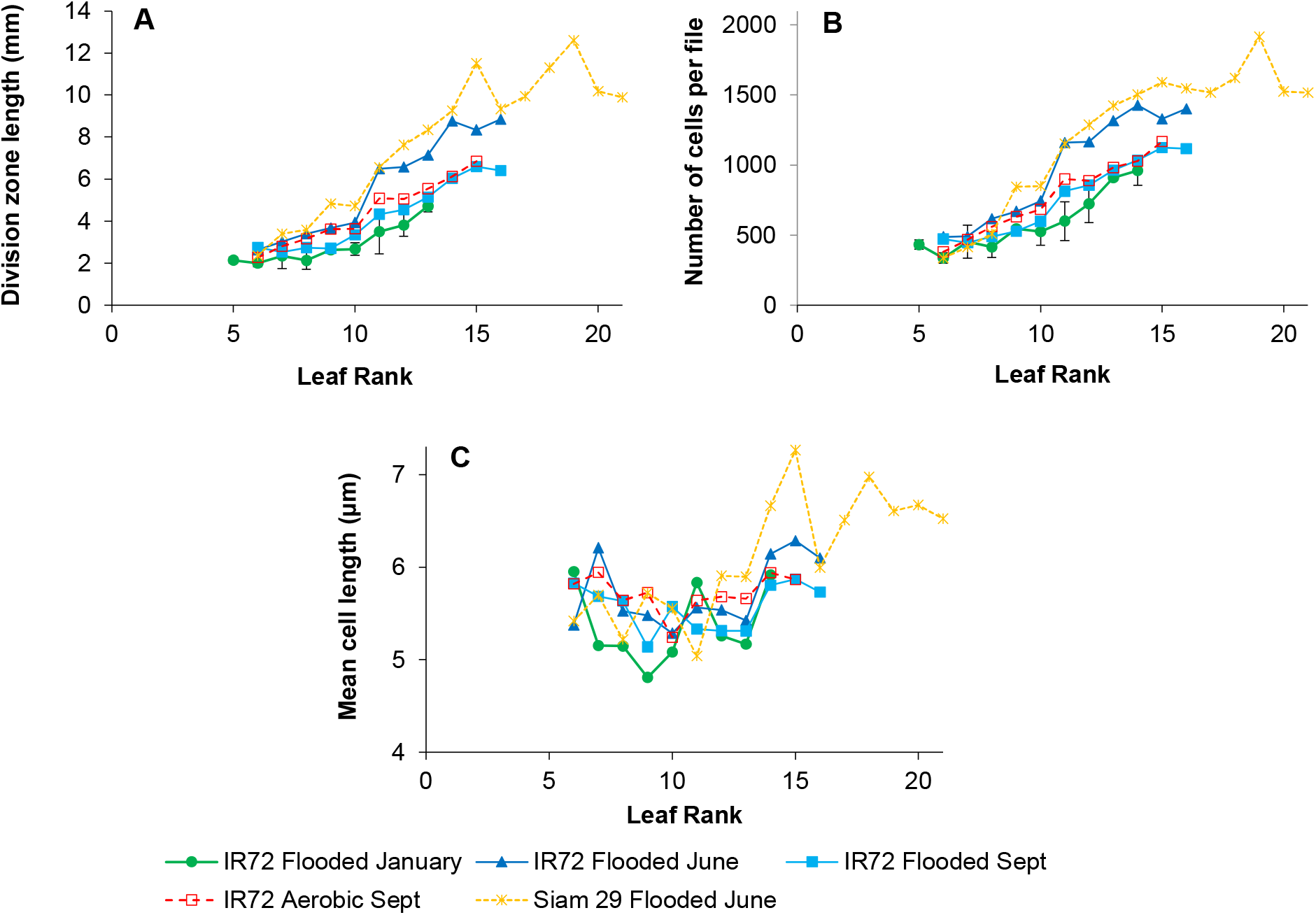
Patterns of the leaf blade meristem division zone with leaf rank on the main stem: length (A) and number of cells (B) of the entire zone, and mean length of the cells (C). Error bars indicate the 95% confidence interval of each mean value from flooded plants of the variety IR 72 sown in January 2015.

The length of leaf blade at sampling (i.e. 3.5- to 4.0-phyllochron-old leaves) did not affect the length and number of cells in the division zone of each leaf rank, except for three Siam 29 leaves (Supporting Information Fig. S9).

### Correlation between leaf blade length and division zone length

The average lengths of leaf blade and meristem division zone of leaves up to rank 11 were strongly correlated (Fig. 4A; R^2^= 0.68, P<0.0001 for IR 72; R^2^= 0.91, P=0.0021 for Siam 29). Similarly, the average length of leaf blade and the average number of cells in the meristem division zone of leaves up to rank 11 were strongly correlated (Fig. 4B; R^2^= 0.70, P<0.0001 for IR 72; R^2^= 0.95, P=0.0006 for Siam 29). However, this linear relationship was lost in leaf 12. From leaf 13, the relationship between leaf blade and division zone lengths plateaued shortly in IR 72 and for the next three leaves in Siam 29; the blade length remained unchanged despite an additional increase in the leaf division zone.

**Fig. 4:**
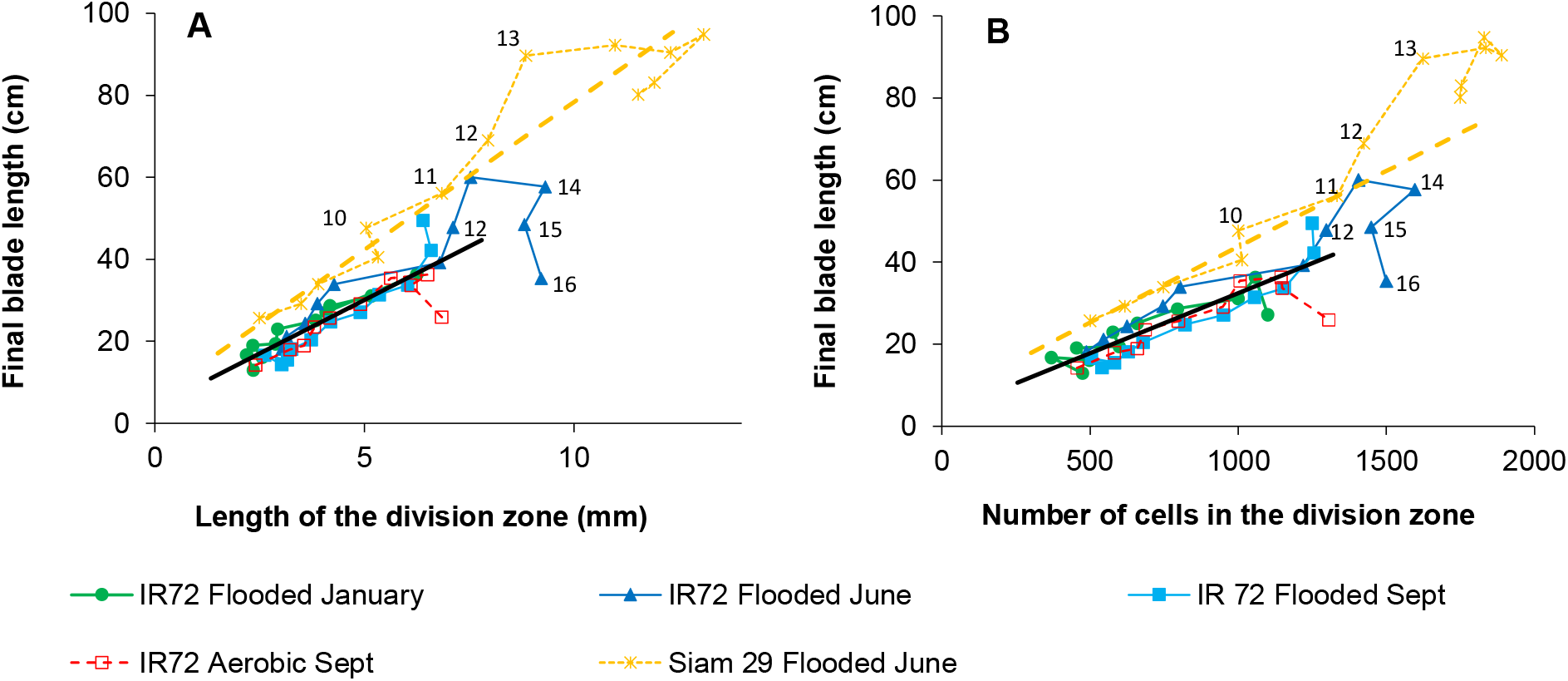
Relationships between the mean length of the blade and mean length (A) or number of cells per line (B) of the leaf meristem division zone in the three experiments. Regressions were plotted for means from leaves of ranks 6 to 11 of IR 72 (straight line) and Siam 29 (dashed line). Rank of some leaves is tagged for the two varieties when sown in June.

### Variations in the mean number of cell doubling cycles in the leaf division zone and their correlation with variations in phyllochron

Homogeneous processes were assumed within the entire division zone and throughout leaf development. The number of cell doubling cycles required to reach the observed cell number per file in the meristem division zone from the initial 1.5 cells in the adaxial epidermis lines increased with leaf rank from 8.5 cycles in leaf 6 to 10.5 cycles in leaf 15 for Siam 29 sown in June, and from 8.3 to 10.1 cycles for IR 72, also sown in June (Fig. 5A). Starting from similar values in leaf 6, the increase in the number of cycles with leaf rank was slower for the other three experimental sets using IR 72, particularly for January sowing, with the maximum number of only 9.3 cycles in leaf 15.

**Fig. 5:**
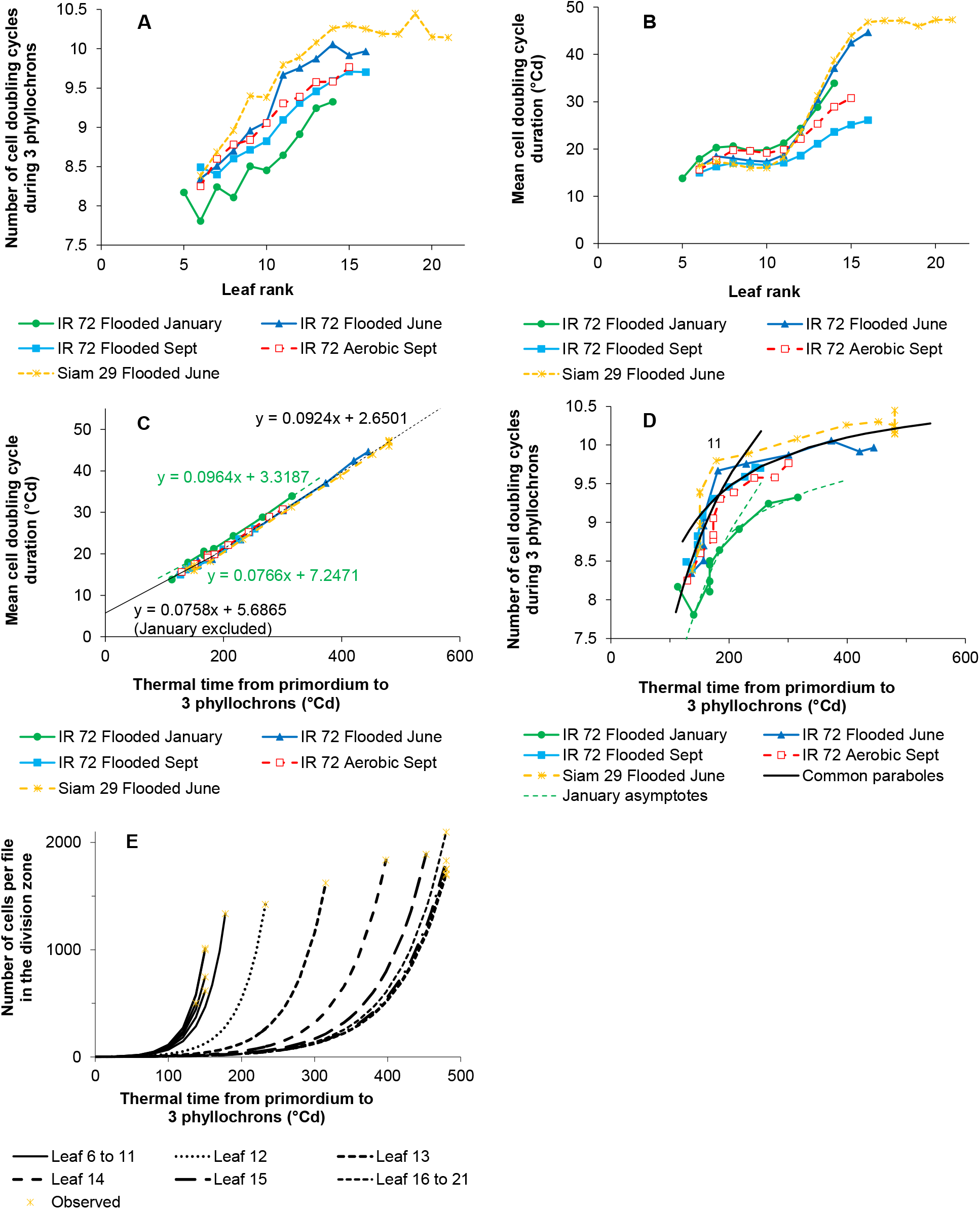
Patterns of cell production in meristems of leaves of rank 5 to 21 during the three experiments: A) Mean number of cell doubling cycles needed to produce the observed maximum number of cells per file in the division zone after three phyllochrons per leaf rank; B) Mean duration of a cell doubling cycle during 3-phyllochrons at each leaf rank; C) Mean number of cell doubling cycles as a function of the 3-phyllochrons duration; D) Relationship between the mean number of cell doubling cycles and the 3-phyllochrons duration ;and E) Predicted dynamics of the cell production during three phyllochrons in the longer files of the division zone of each leaf of the variety Siam 29 sown in June.

The mean doubling cycle duration was rather stable from leaves 6 to 11, followed by a sharp increased from leaves 12 to 16 (Figs. 5B and 1A). From leaf 12 onwards, the mean doubling cycle duration remained lower in plants sown under flooded conditions in September than in plants subjected to the other treatments. From leaves 6 to 11, the mean doubling cycle duration was longer for IR 72 sown under flooded conditions in January than for the other varieties subjected to the remaining treatments. The durations of mean doubling cycle and three phyllochrons in the four June and September experiments showed a strong linear relationship (R^2^=0.9973, P<0.0001), albeit with an uneven distribution of residuals (Fig. 5C). Therefore, the linear relationship were separately computed for data until leaf 11 (R^2^=0.92, P<0.0001) and for data from leaf 12 onwards (R^2^=0.998, P<0.0001). The same relationships were also computed for January sowing.

The doubling cycle duration was *t*_×*2*_ *= a + b* × *P*_*3*_. If the number of doubling cycles (N) during *P*_*3*_ is 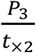, then 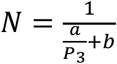, which is an asymptotic function of *P*_*3*_, with the limit of 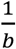, as shown in Fig. 5D.

The simulated dynamics of cell production in the division zone with thermal duration were plotted by leaf for Siam 29 sown in June using equation 4 (Fig. 5E).

In IR 72, the plastochron was stable (48.9 °Cd) for leaves 7–10 whereas the number of cells per file in the division zone increased (Supporting Information Table S1). The phyllochron slowly increased with leaf rank (6.5% per leaf) because of the increasing length of the sheaths until leaf 13 (Clerget et al., 2014) that delayed the appearance of the leaf tips, which in turn delayed the triggering of the formative divisions (Supporting Information Fig. S10). Assuming a stable rate of cell production in these four leaves, small increases of the duration of the proliferative division phase caused exponential increases of the division zone length. Beyond leaf 13, sheath length (data not shown) and phyllochron were stable in Siam 29.

## DISCUSSION

### Broken-linear models well fit the leaf appearance kinetics that were similar for both varieties but sensitive to the sowing date

The 2-day interval between records of the number of leaves on the main stem of the two varieties directly sown in June in a puddle field clearly showed that leaves appeared linearly with time, albeit at rates that decreased twice or thrice during the plant lifetime and with smoothed transition phases between two developmental rates (Supporting Information Fig. S7). These field observations corroborate the results of previous greenhouse-based experiments (Egle *et al*., 2015). A crucial consequence of these discontinuities in the rate of leaf initiation and appearance was the discontinuous increase in the duration of leaf development with leaf rank (Fig. 1B).

The faster rate of leaf appearance observed in flooded plants of IR 72 sown in September was not anticipated because of the unusual date. This was possibly a photoperiod-sensitive reaction similar to that observed previously in sorghum sown before the autumn equinox (Clerget *et al*., 2008). Therefore, differences in sowing dates must be carefully analysed in the light of this crucial factor for plant development.

### Reason the observed division zone was shorter than that reported previously

The mean length of the division zone of leaf 6 was close to 2.5 mm in IR 72 and Siam 29 (Fig. 3A), compared with 14 mm in the *indica* variety IR 64 and 20 mm in the tropical *japonica* variety Azucena, both for leaf 6, reported by Parent *et al*. (2009) and 5.8 mm for leaf 3 in the japonica variety Chunjiang 06 reported by Fang *et al*. (2018). Similarly, the mean dividing cell length in leaf 6 was 5.4 μm in IR 72 and Siam 29 (Supporting Information Fig. S8), compared with 8.3 μm in IR 64 and 10 μm in Azucena for leaf 6 (Parent *et al*., 2009) and 13 μm in Chunjiang 06 for leaf 3 (Fang *et al*., 2018). Furthermore, the mean number of cells per file of division zone for leaf 6 was 487 in IR 72 and 337 in Siam 29, compared with 1,687 in IR 64 and 2,000 in Azucena for leaf 6 (Parent *et al*., 2009) and 491 in Chunjiang 06 for leaf 3 (Fang *et al*., 2018). In the present experiment, the division zone was restricted to the proliferative division zone, consistent with the observations reported by Fiorani *et al*. (2000). Conversely, in the cited previous studies, the division zone extended until the end of formative divisions. Here, in leaf 10 of IR 72, the proliferative zone ended at 4.1 mm, whereas the formative divisions of guard cells were not observed until 7 mm (Supporting Information Fig. S4), indicating that the end of the formative division zone was more distal than 7 mm. Consistent with the observations of Fiorani *et al*. (2000) and Masle (2000), no proliferative division with straight immature cell walls was observed beyond 4.1 mm. Thus, the first asymmetrical divisions producing half-moon-shaped cell walls indicate the frontier between the proliferative and formative zones of cell divisions. This modified definition led to a shorter division zone than that reported previously.

### Division zone length profile shaped the leaf blade length profile

In all five experiments, the length of the meristem cell division zone and the length of the blade were linearly correlated up to leaf 11; however, the relationship was variety-dependent (Fig. 4A). Such a linear relationship between these two lengths has been reported in wheat (Kemp, 1980) and *Poa* (Fiorani *et al*., 2000). The present experiment extended the examination to all leaves on the stems of rice plants, but concluded that the correlation changed after leaf 11. Logically, longer meristematic cell files would generate more cells entering the elongation zone during the 1.5 phyllochrons dedicated to blade development. The shift in the relationship between the division zone and blade lengths for leaves 12 and 13 of June-sown varieties occurred simultaneously with a large increase in phyllochron (Figs. 1 and 4). In IR 72, the division zone length of the penultimate leaf 15 and flag-leaf 16 was the maximum, although their blade lengths were much shorter than those of the previous leaves. These two leaves develop and grow simultaneously with the ear, and either they compete for nutrition and space or the development of their blades is down-regulated. In Siam 29, the vegetative phase continued until the end of the experiment, and a substantially stable relationship between the division zone and blade lengths was maintained from leaf 14 onwards.

### Linkages amongst the cell production rate, plastochron, and phyllochron

Rice is a unique cereal, in which the plastochron and phyllochron are equal throughout the vegetative phase (Nemoto *et al*., 1995). Strong linear correlations between the three phyllochron durations of the leaf proliferative division zone growth and the mean rate of cell production in the meristems of leaves 6–21 are shown in Fig. 5C. The correlation changed between leaves 11 and 12, simultaneously with changes in the leaf appearance rate (Fig. 1 and Supporting Information Fig. S7) and the linear relationship between the length of the division zone and final blade (Fig. 4). The link between the rate of cell division in the stem apex and plastochron has been previously reported (Nougarède *et al*., 1990; Itoh *et al*., 1998; Cockcroft *et al*., 2000; Kawakatsu *et al*., 2006). Accordingly, in rice, the rate of apical cell division decreased in steps at the primordium initiation of leaves 5, 12, and 21 whilst remaining stable during the intervals. These decreases plausibly reduced the growth rate of the stem apical meristem and consequently increased the duration between the initiation of two successive primordia. Indeed, the initiation of leaf primordia is controlled by hormonal gradients and occurs at a stable distance from the apical tip (Xue *et al*., 2020). The rate of cell division is higher in the leaf meristem, which rapidly encloses the stem apex, than in the apical meristem; however, the relationship between these two rates remains unknown. However, the linear correlation between the rate of cell production in the leaf meristems and phyllochron leads to the conclusion that the rates of cell production in both apical and axillary leaf meristems were linked through stable coefficients that only changed when the rate of cell production in the apical meristem changed.

Equality of the plastochron and phyllochron was not respected in leaves 7–10 while the sheath length was increasing from one leaf to the next (Supporting Information Table S1). In agreement with Skinner & Nelson (1995), the phyllochron increased slowly during this phase because of the resulting delayed tip appearance (Supporting Information Fig. S10). The total increase was only 20%, thus difficult to detect before the next breaking point in the appearance kinetics. In contrast, the lengths of the division zone exponentially increased leaf after leaf in response to the small increases of the phyllochron, which resulted in significant increases of the blade length.

In summary, the vertical leaf length profile in rice plants of a specific variety growing under a given favourable environment is determined by the rate of cell division in the stem apical meristem, which also determined the plastochron and phyllochron duration. Leaf sheaths are generated by the same division zone and, consequently, follow the same patterns that are also expected for internode cell production.

### Effect of experimental factors on the leaf size profiles

#### Varieties

The leaf area profile of IR 72 sown in June under flooded conditions, with the maximum area of 57 cm^2^ and the maximum length of 60 cm for leaf 14, displayed the highest values ever observed on the IRRI farm for this variety (unpublished data) (Fig. 2C). In Tokyo, Okami *et al*. (2012) reported an area of only 40–45 cm^2^ for leaf 14 of IR 72 grown in the field, whereas Egle *et al*. (2015) reported the maximum length of 60 cm for leaf 14 of IR 72 in pots installed in a greenhouse.

For each leaf rank, the leaf blades of Siam 29 were much longer than those of IR 72 sown in June, with the maximum lengths of respectively 95 and 60 cm (Fig. 3A). In contrast, the length and number of cells in the division zones of all leaves in Siam 29 were only slightly higher than those in IR 72 (Fig. 4A&B and Table 2). IR 72 is an improved semi-dwarf variety bearing the *sd1* dwarfing gene, whereas Siam 29 is a tall landrace. The *sd1* gene reduces gibberellic acid concentration and vegetative organ size (Sasaki *et al*., 2002). The *sd1* gene dwarfs stem length mainly through decrease in the number of cells in the internode cell files (Kamijima, 1991; Ogi *et al*., 1993), although no information is available on leaf cells. Assuming that *sd1* produces the same effect in leaf blades as in the internode, it only occurred during the P3-P4.5 phase when the rate of cell division was maximum in the proliferative division zone (Parent *et al*., 2009). Indeed, the difference between the mean division zone lengths was only 14% (Fig. 3A&B), and the relationship between the lengths of the division zone and leaf blade differed between the two varieties (Fig. 4). This difference likely resulted from the 50% increase in the final cell length in Siam 29 compared with that in IR 72, which was not measured in the present experiment.

#### Sowing date

IR 72 was sown and managed under flooded conditions in January, June, and September. In plants sown in June, the lengths of leaf blade and leaf division zone were much higher than those in the other months (Figs. 2A and 3A&B). In plants sown in September, the leaf appearance rate was much faster than that in the other months (Fig. 1A), resulting in the shorter duration of division zone growth (Fig. 1B) and shorter leaf blades. In plants sown in January, leaf blades were shorter because of the lower rate of cell division in leaf meristems than in the other months. The setup probably occurred during seedling emergence, when young plants sense the environment and acclimate accordingly. Similar responses of phyllochron and duration to flowering initiation to temperature or photoperiod have been reported (Kirby, 1995; Birch *et al*., 1998; Clerget *et al*., 2021).

#### Watering management

In September, IR 72 was either kept flooded with a 5 cm water layer or grown in aerobic soil frequently re-irrigated to maintain soil moisture over the field capacity. As previously reported, the leaf appearance rate is slower in aerobic than in flooded environments (Clerget *et al*., 2014). In the present study, the leaf and division zone lengths were higher in the aerobic environment (Figs. 3 and 4), which is inconsistent with previous observations. However, similar observations have been recorded in a previous greenhouse experiment in which IR 72 plants were grown under flooded and aerobic environments. Leaf appearance was faster (Clerget & Bueno, 2013) but mean leaf length above rank 10 was significantly lower (P=0.02) in flooded pots than in aerobic pots (unpublished data). Thus, in both experiments, the leaves of directly seeded plants developed faster under flooded conditions and bore shorter division zones and leaf blades, corroborating the model presented in the present and previous studies (Rebetzke & Richards, 1999; Kawakatsu *et al*., 2006; Rebolledo *et al*., 2012). Furthermore, the greater length of leaves in transplanted flooded plants than in aerobic direct-seeded plants may be attributed to the nursery effect. The developmental delay experienced in the nursery accelerates the rate of development after transplanting (Clerget *et al*., 2021). The same effect would also modify the ratio between the rate of cell division in the apical meristem and leaf meristems, as observed in January sowing. Consequently, transplanted flooded plants developed faster than aerobic plants, whilst their leaf meristems produced cells at an accelerated rate to counterbalance the usual effect of the faster rate of leaf appearance on leaf size. Thus, the millenaries-old Asian practice of rice transplanting takes advantage of two highly efficient underlying mechanisms of plant acclimation.

## Supporting information

Supplementary Pablo 2022 Rice leaf profile

## ACKNOWLEDGEMENTS

We thank Dr Paul Quick who supported the project and gave access to the imaging facilities of the IRRI-C4 laboratory; Pedro Gapas, Rene Carandang, Luis Malabayabas, and Victor Lubignan who helped with the crop physiology experimental tasks; and Aya Mercado for the microscopy works.

## Supporting information

Additional Supporting Information may be found online in the Supplementary Material section.

**Fig. S1:** Measurement of the length of a next-to-appear leaf before cutting off 10-15 mm at the leaf basis for meristem cell imaging.

**Fig. S2**: In the division zone, three types of cell files constitute the epidermis. A: Leaf base transversal section and B: photomicrograph of the adaxial leaf epidermis in the division zone.

**Fig. S3**: Distally oriented half-moon shaped asymmetric divisions clearly showing the distal end of the division zone.

**Fig. S4:** Cell length profiles in one stomatal cell file of the blade meristem of one leaf 10 of the variety IR 72 sown in January 2015 and 100-μm long pictures of the cell files at **A)** the first occurrence of a formative asymmetrical division, **B)** 50% occurrence of the asymmetrical division, **C)** completion of the asymmetrical divisions, and **D)** further cell differentiation.

**Fig. S5**: Apical initiation (A) and chronological development of leaf 11 during the first phyllochron (B, C) in the main stem of rice plants. Courtesy of M. de Raissac and J.L. Verdeil, 2006, CIRAD.

**Fig. S6:** Monthly means of A) daily mean, maximum and minimum temperatures and B) daily global solar radiation for the three experiments under flooded conditions sown in January, June and September 2015.

**Fig. S7:** Mean observed and predicted number of appeared leaves with thermal time in A. IR 72 and B. Siam 29 sown in June 2015.

**Fig. S8:** Samples of the cell length profiles in the leaf blade division zones of the varieties IR 72 and Siam 29 sown in June 2015.

**Fig. S9:** Cell number in the intercostal cell lines of the division zone plotted versus the blade length at the sampling time of each sampled leaf for each leaf rank.

**Fig. S10:** Estimated length of leaves 7–10 from initiation to tip appearance. The dashed arrow represents the cue that trigger the onset of the formative divisions in the division zone at the tip appearance of the previous leaf.

**Table S1:** Optimized plastochron and phyllochron3 values in leaf 7–10 of IR 72 sown in June to fit the observed number of cells per file in the division zone at P3 and the calculated phyllochron of 52°Cd.

