## Supplementary Pablo 2022 Rice leaf profile for "Phyllochron duration and changes through rice development shape the vertical leaf size profile"

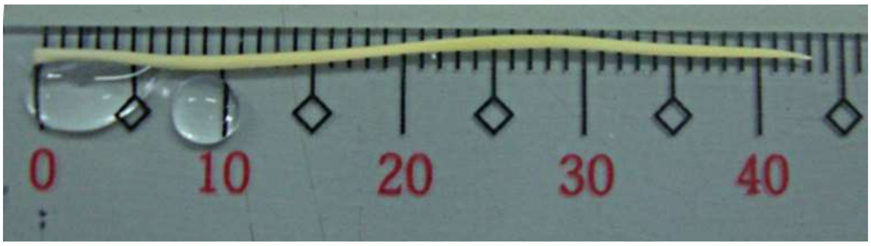


**Fig. S1:** Measurement of the length of a next-to-appear leaf before cutting off 10-15 mm at the leaf basis for meristem cell imaging.

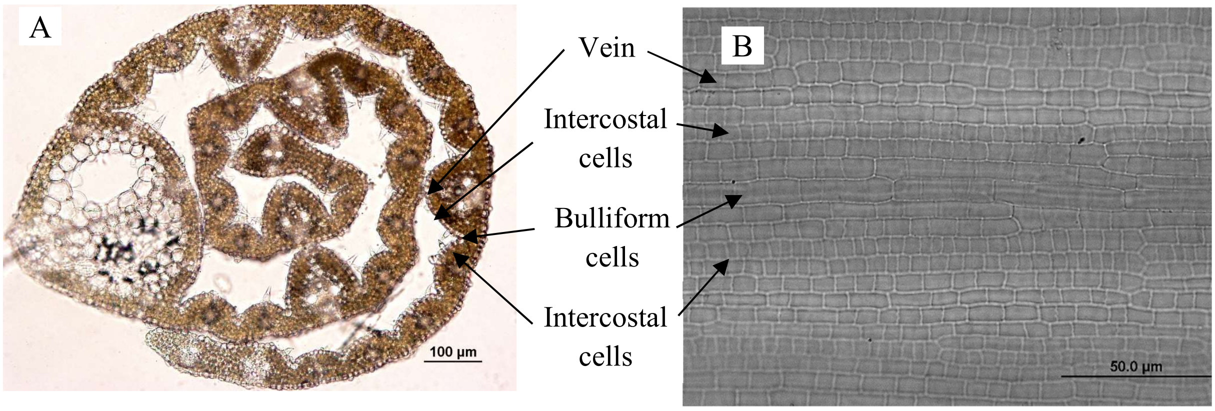


**Fig. S2:** In the division zone, three types of cell files constitute the epidermis. A: Leaf base transversal section, and B: photomicrograph of the adaxial leaf epidermis in the division zone

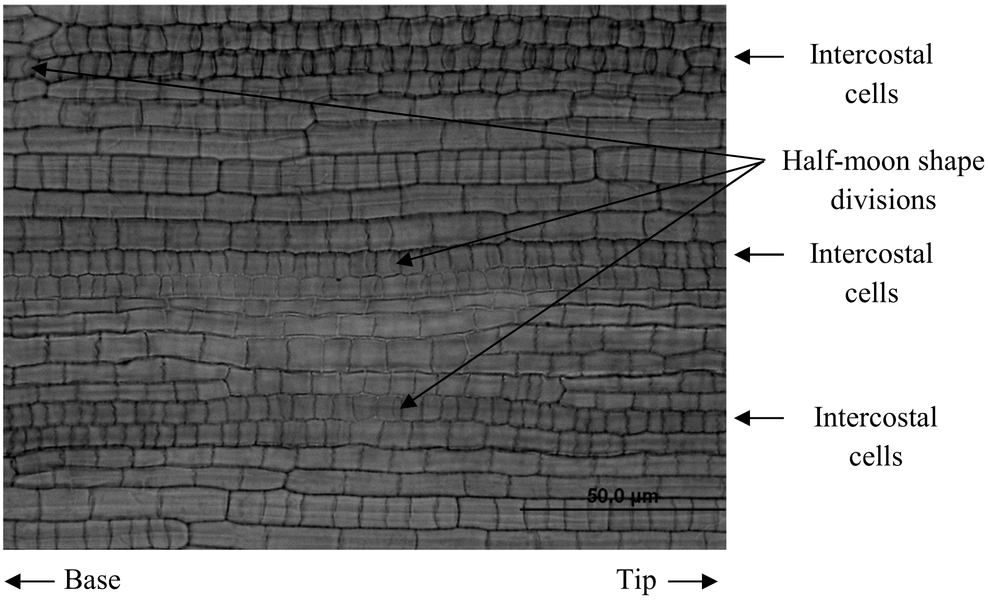


**Fig. S3:** Distally oriented half-moon shaped asymmetric divisions clearly showing the distal end of the division zone.

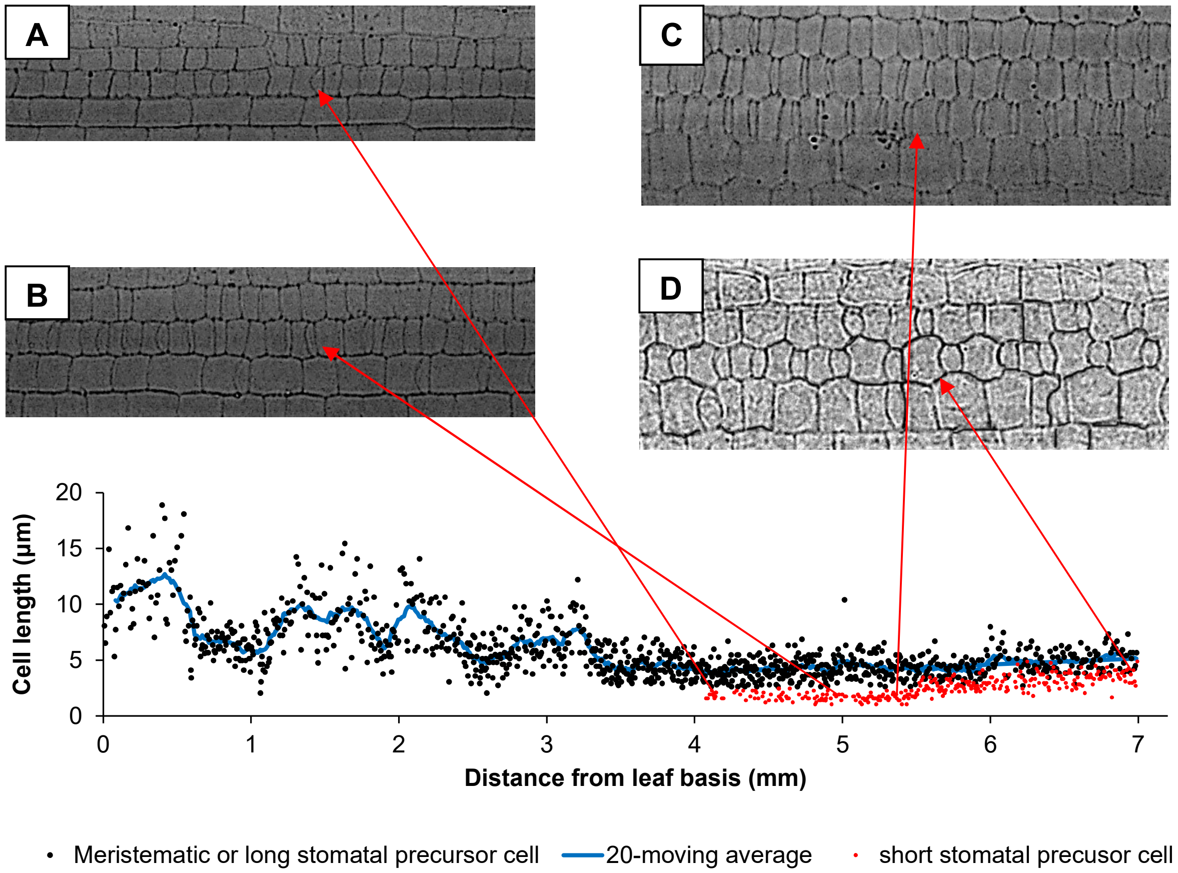


**Fig. S4:** Cell length profiles in one stomatal cell file of the blade meristem of one leaf 10 of the variety IR 72 sown in January 2015 and 100-µm long pictures of the cell files at **A)** the first occurrence of a formative asymmetrical division, **B)** 50% occurrence of the asymmetrical division, **C)** completion of the asymmetrical divisions, and **D)** further cell differentiation.

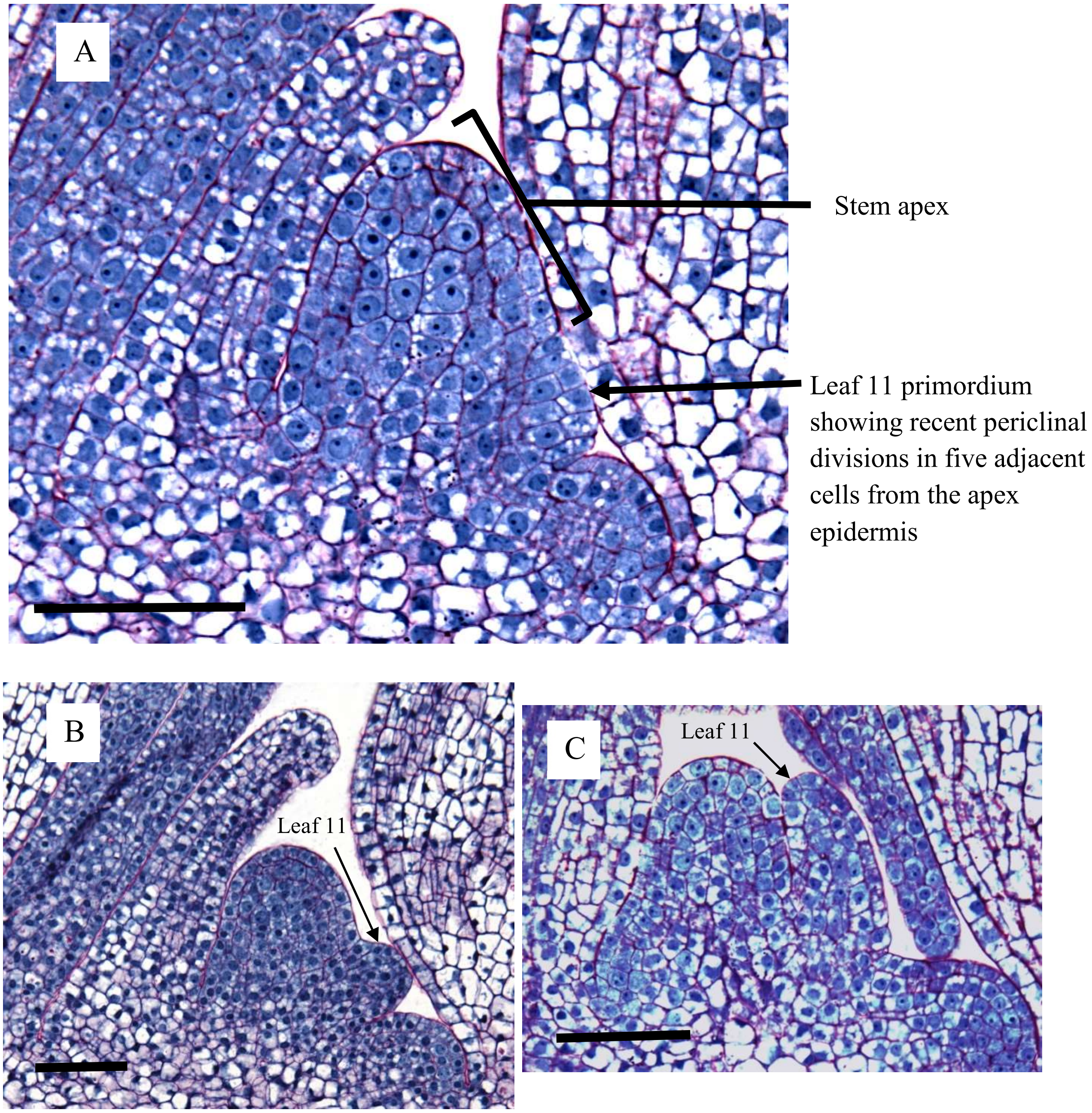


**Fig. S5:** Apical initiation (A) and chronological development of leaf 11 during the first phyllochron (B,C) in the main stem of rice plants. Bars in the left lower corners are 50 µm long. Courtesy of M. de Raissac and J.L. Verdeil, 2006, CIRAD, France.

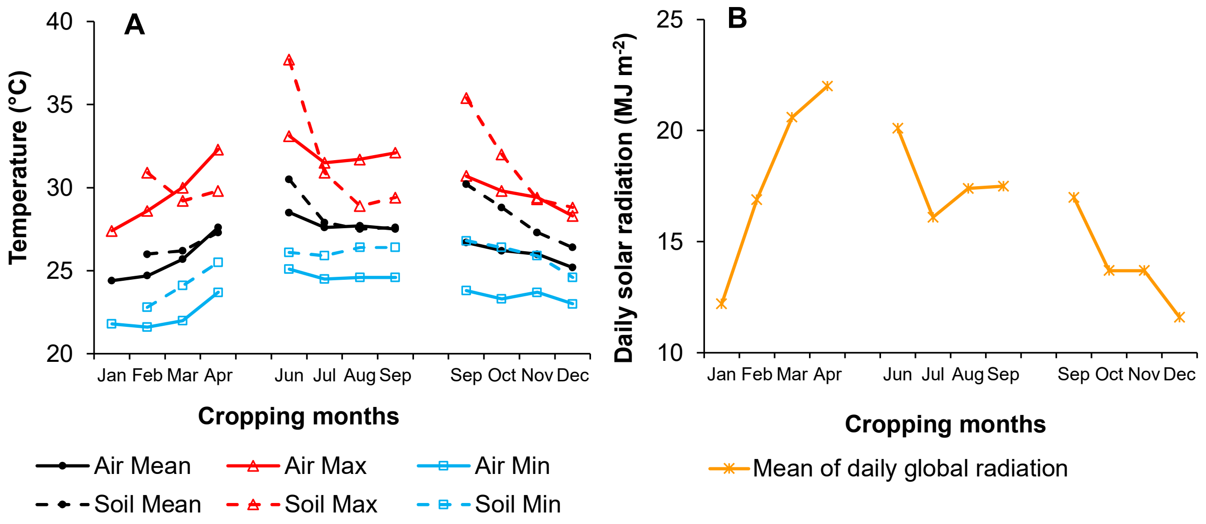


**Fig. S6:** Monthly means of A) daily mean, maximum and minimum temperatures and B) daily global solar radiation for the three experiments under flooded conditions sown in January, June and September 2015.

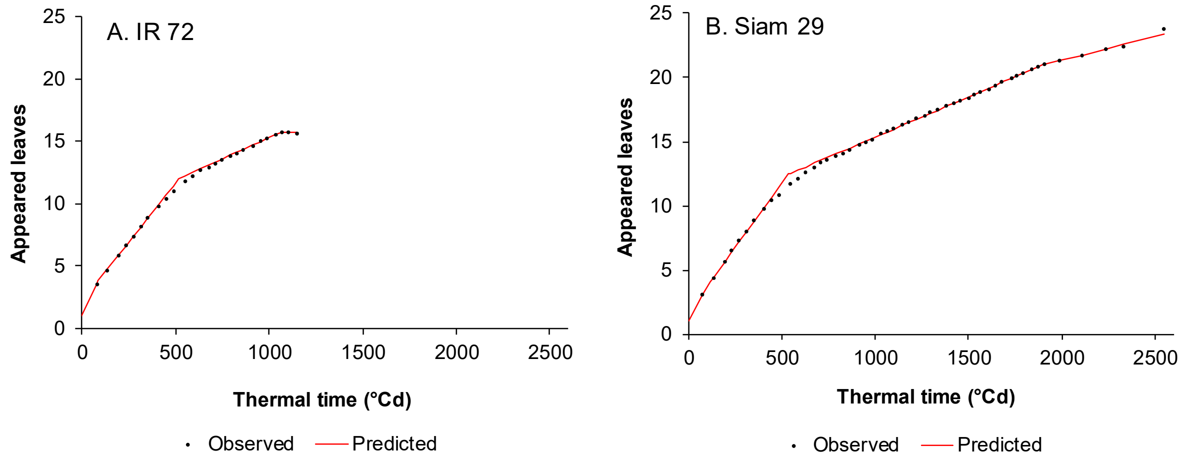


**Fig. S7:** Mean observed and predicted number of appeared leaves with thermal time for A. IR 72 and B. Siam 29 sown in June 2015.

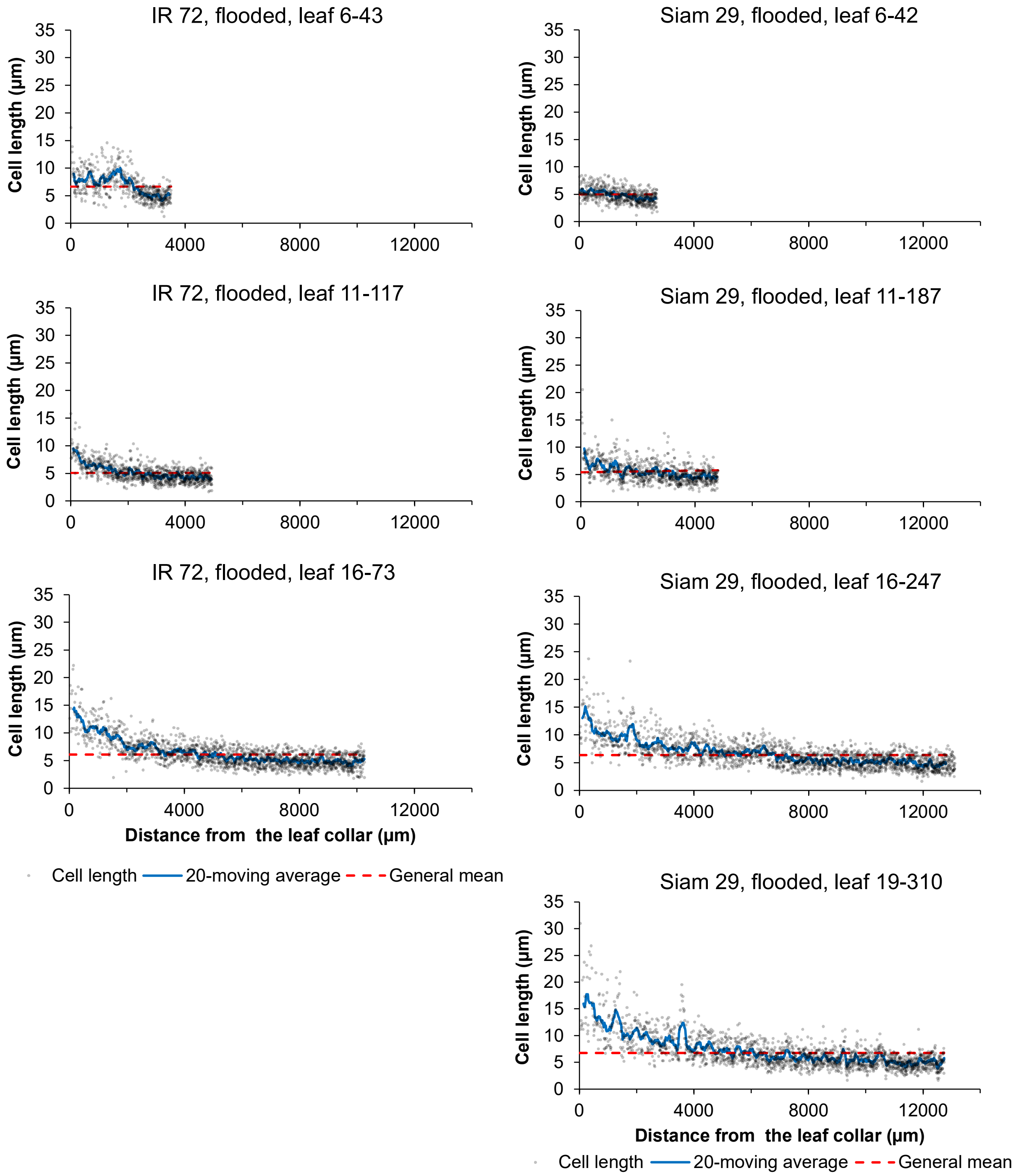


**Fig. S8:** Samples of the cell length profiles in leaf blade division zones of the varieties IR 72 and Siam 29 sown in June 2015. Leaves were numbered with their rank and their length at sampling date.

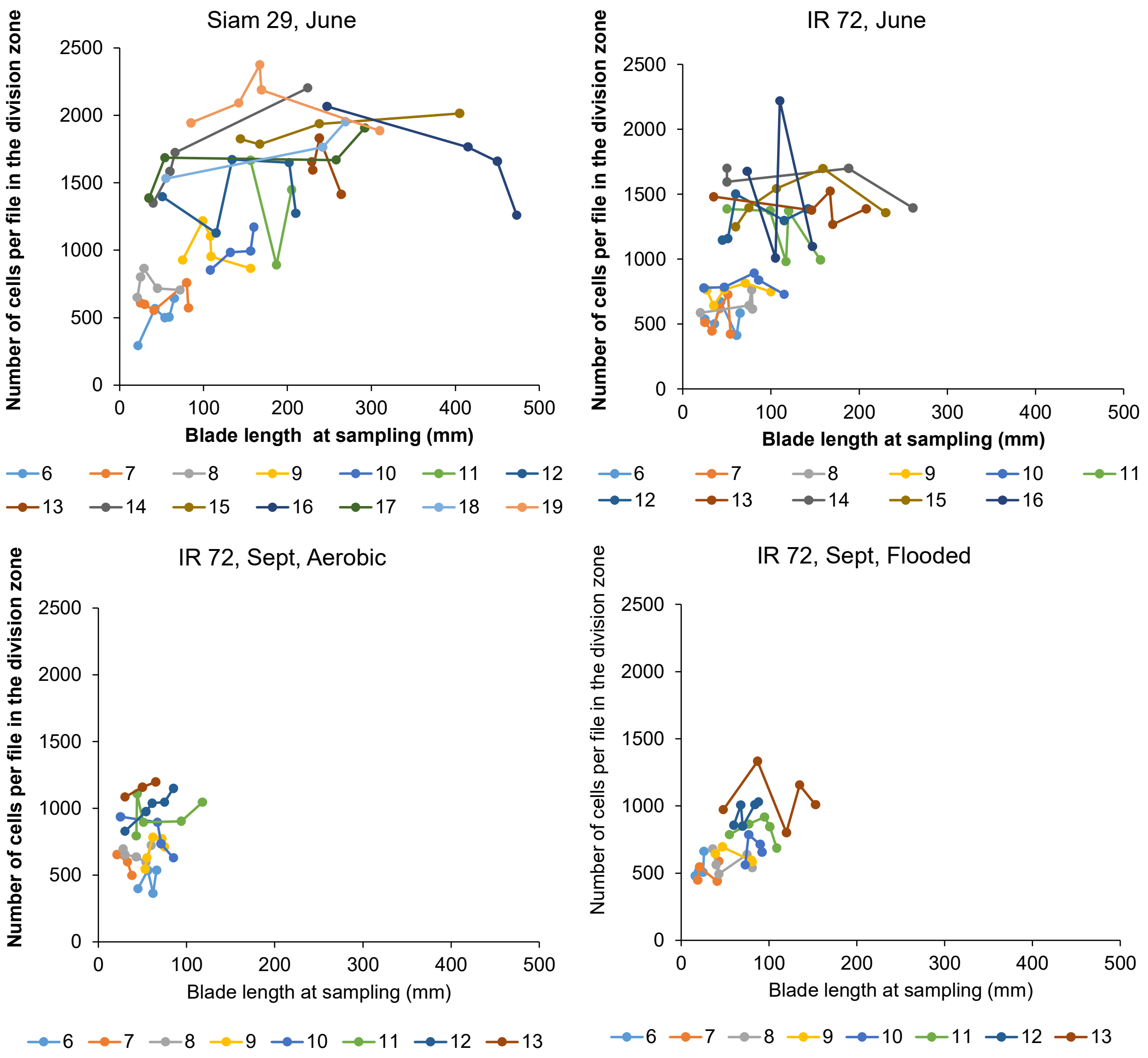


**Fig. S9:** Cell number in the intercostal cell lines of the division zone plotted versus the blade length at sampling time for each sampled leaf and for each leaf rank.

**Table S1:** Optimized plastochron and phyllochron3 values in leaf 7–10 of IR 72 sown in June to fit the observed number of cells per file in the division zone at P3 and the calculated phyllochron of 52°Cd.

| Leaf | plastochron | phyllochron3 | P3 | Tdiv | Number of cells at P3 | | |
|---|---|---|---|---|---|---|---|
| | °Cd | °Cd | °Cd | °Cd | observed | predicted | Square error |
| 7 | 48.9 | 46.6 | 144 | 16.9 | 546 | 546.02 | 0.00062 |
| 8 | 48.9 | 49.9 | 147 | 16.9 | 625 | 624.98 | 0.00061 |
| 9 | 48.9 | 54.2 | 152 | 16.9 | 746 | 745.97 | 0.00082 |
| 10 | 48.9 | 56.1 | 154 | 16.9 | 804 | 804.03 | 0.00087 |

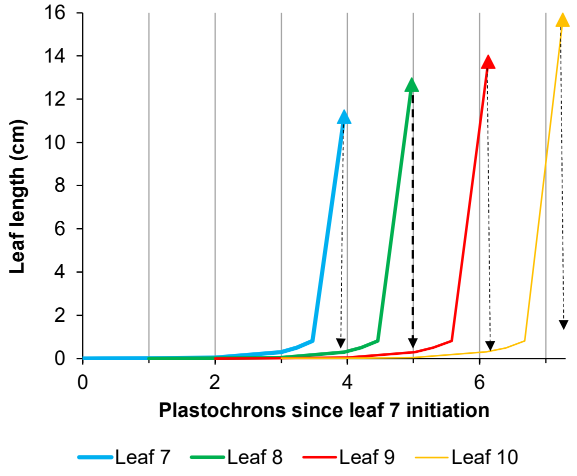


**Fig. S10:** Estimated length of leaves 7–10 from initiation to tip appearance. The dashed arrow represents the cue that trigger the onset of the formative divisions in the division zone at the tip appearance of the previous leaf.
